# Tensor decomposition–based unsupervised feature extraction for integrated analysis of TCGA data on microRNA expression and promoter methylation of genes in ovarian cancer

**DOI:** 10.1101/380071

**Authors:** Y-h. Taguchi, Ka-Lok Ng

**Affiliations:** Department of Physics Chuo University, 1-13-27 Kasuga Bunky-ku, Tokyo 112-8551, Japan; Department of Bioinformatics and Medical Engineering Asia University, Taichung, Taiwan, Department of Medical Research, China Medical University Hospital China Medical University, Taiwan

**Keywords:** tensor decomposition, feature extraction, ovarian cancer, microRNA, promoter methylation

## Abstract

Integrated analysis of epigenetic profiles is important but difficult. Tensor decomposition–based unsupervised feature extraction was applied here to data on microRNA (miRNA) expression and promoter methylation of genes in ovarian cancer. It selected seven miRNAs and 241 genes by expression levels and promoter methylation degrees, respectively, such that they showed differences between eight normal ovarian tissue samples and 569 tumor samples. The expression levels of the seven miRNAs and the degrees of promoter methylation of the 241 genes also correlated significantly. Conventional Student’s *t* test–based feature selection failed to identify miRNAs and genes that have the above properties. On the other hand, biological evaluation of the seven identified miRNAs and 241 identified genes suggests that they are strongly related to cancer as expected.

## I. INTRODUCTION

Multiomics analysis is key to understanding complicated regulation of gene expression by multiple factors. Examples of these factors are DNA methylation [1], histone modification [2], and chromatin structures [3]. Functional noncoding RNAs, including microRNA (miRNA) [4], are also often regarded as some of important regulators of gene expression. In spite of the importance of these regulators, it is rarely discussed how multiple factors regulate gene expression cooperatively.

Especially, the relation between methylation and miRNA as regulators is unclear, although how methylation affects miRNA expression is discussed [5]. The reason why this topic is not discussed much is possibly that methylation contributes to pretranscriptional regulation whereas miRNA contributes to post-transcriptional regulation. Because it is difficult to figure out how methylation and miRNA can regulate gene expression cooperatively from the biological point of view, data-driven approaches are the only possible strategy. To this end, we need to identify a set of genes to which the amount of methylation is attributed and miRNAs that fulfill the following conditions.

1. MiRNA should be expressed differentially between treated and control samples.
2. The degrees of promoter methylation of the genes should be different between treated and control samples.
3. Expression levels of these miRNAs and the degrees of promoter methylation of the above genes should significantly correlate.

If a set of miRNAs and genes fulfills these criteria, then they are candidates that regulate gene expression cooperatively, although further analysis will be necessary to see if they really work cooperatively. The purpose of this study was restricted to the identification of a set of miRNAs and genes that satisfy the above three conditions.

Nonetheless, even finding the sets of miRNAs and genes that fulfill these *weak* conditions (by expression levels and degrees of promoter methylation, respectively) is not easy. First of all, because we cannot restrict pairing of miRNA expression levels and degrees of promoter methylation of genes, all possible pairs must be tested. The number of possible pairs can easily exceed a few million. This means that *P*-values must be at least smaller than 1 × 10^−6^ × 0.05 ≃ 1 × 10^−8^ if the required possible threshold *P*-value is 0.05. It is generally not easy to identify such highly significant pairs of differentially expressed miRNAs and differentially methylated genes. Second, miRNAs and genes showing differences in expression levels between treated and control samples do not always correlate. If the difference between controls and treated samples is not large enough, the correlation between expression of miRNAs and promoter methylation data on genes is not governed by the differences between control and treated samples but rather by correlation within each class: miRNA expression vs gene methylation in normal tissues or miRNA expression vs gene methylation in tumors. For example, The Cancer Genome Atlas [6] (TCGA) sample is not associated with equal numbers of treated and control samples but is associated with a combination of very small numbers of control samples and large numbers of tumor samples; this means that the correlation between miRNA expression levels and promoter methylation degrees of genes is governed by that within tumor samples.

To overcome this difficulty, we employed tensor decomposition (TD)-based unsupervised feature extraction (FE) [7], [8], [9], [10], [11], [12], [13]. TD-based unsupervised FE was applied to ovarian methylation profiles and miRNA expression data retrieved from TCGA. We successfully obtained a set of methylation sites and miRNAs that significantly correlate and show a significant dissimilarity between controls and treated samples simultaneously. Enrichment analysis also identified biological significance of the differentially expressed miRNAs and differentially methylated genes.

## II. MATERIALS AND METHODS

### A. Methylation profiles and miRNA expression

Ovarian methylation profiles and miRNA expression data were downloaded from TCGA. They are composed of eight normal ovarian tissue samples and 569 tumor samples. Our dataset includes expression data on 723 miRNAs as well as promoter methylation profiles of 24906 genes.

### B. TD-based unsupervised FE

Given that the method was described in detail in another paper [10], it is described here only briefly. Suppose that 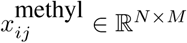 is the degree of promoter methylation of the *i*th gene of the *j*th sample whereas 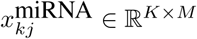 is the expression level of the *k*th miRNA of the *j*th sample. *N* (= 24906) is the number of genes whose promoter methylation status is known, and *K* (= 723) is the number of miRNAs whose expression has been measured, and *M* (= 577) is the number of samples. Both *x*_*ij*_ and *x*_*kj*_ were standardized such that they were associated with zero mean and unit variance, i.e., 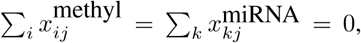 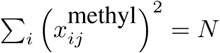, and 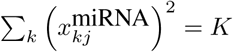.

Next, to generate a case II type I tensor [10], we define

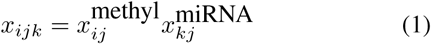

*x*_*ijk*_ was subjected to Tucker decomposition as follows:

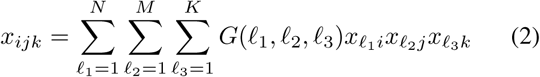

where *G* is the core tensor and *x*_*l*_1_*i*_ ∈ ℝ^*N* × *N*^, *x*_*l*_2_*i*_ ∈ ℝ^*M* × *M*^, *x*_*l*_3_*i*_ ∈ ℝ^*K* × *K*^are singular value matrices that are orthogonal. Because Tucker decomposition is not unique, we have to specify how we derive Tucker decomposition. In particular, we chose higher-order singular value decomposition (HOSVD) [14].

Given that *x*_*ijk*_ is too large to apply TD as is, we generate a case II type II tensor

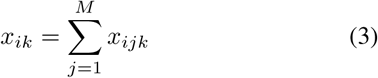

Singular value decomposition (SVD) was applied to matrix *X* ∈ ℝ^*N* × *K*^ whose components are *X*_*ij*_; thus, we get

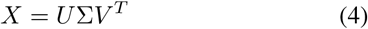

where *U* ∈ ℝ^*N* × *K*^ and *V* ∈ ℝ^*K* × *K*^ are orthogonal matrices (here *N* > *K*), and ∑ ∈ ℝ^*K* × *K*^ is a diagonal matrix. *U*^*T*^ should correspond to *x*_*l*_1_*i*_. This means that *x*_*l*_1_*i*_ = 0 for *l*_1_ > *K*. On the other hand, *V*^*T*^ should corresponds to *x*_*l*_3_*k*_.

*x*_*l*_2_*j*_ that corresponds to samples cannot be obtained by SVD. As shown in the previous study [10], we can obtain two *x*_*l*_2_*j*_s that correspond to methylation and miRNA, respectively, in the following way:

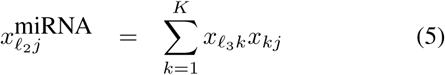

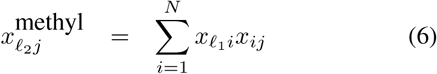

The selection of genes to which methylation profiles are attributed and miRNAs using the above results can be performed as follows. First, among singular value vectors attributed to samples, we select 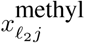 and 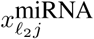 that show significant differences between normal ovarian tissues (1 ≤ *j* ≤ 8) and tumors (*j* > 8). This task can be accomplished, for example, with some statistical tests like Student’s *t* test. Suppose that 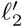 turned out to show dissimilarity between control and treated samples. Then, *P-*values are attributed to *k* miRNAs and *i* genes, assuming that *x*_*l*_1_*i*_ and *x*_*l*_3_*k*_ obey a normal distribution,

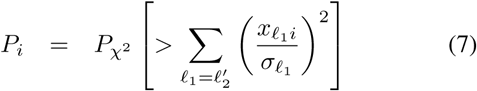

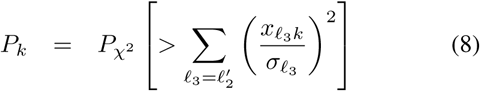

where *P*_*χ*^2^_ [> *x*] is cumulative probability that the argument is greater than *x* in a *χ*^2^ distribution. *σ*_*l*_1__ and *σ*_*l*_3__ are standard deviations for *x*_*l*_1_*i*_ and *x*_*l*_3_*k*_, respectively. After *P-*values are adjusted by means of the Benjamini-Hochberg (BH) criterion [15], miRNAs and genes that are associated with adjusted P-values less than 0.01 are selected as those showing differences in expression and promoter methylation, respectively, between controls (normal ovarian tissues) and treated samples (tumors).

### C. Analysis of correlation between expression of the miRNAs and promoter methylation of the genes

Pearson’s correlation coefficients were computed by the `corr` function in R software. *P*-vales were computed with the `cor.test` function of R.

## III. RESULTS

We applied TD-based unsupervised FE to an ovarian cancer dataset retrieved from TCGA. Then, we found that 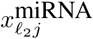 and 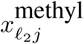 with *l*_2_ = 2 are different between normal ovarian tissues and tumors. Student’s *t* test was performed on the two sets, *j* ≤ 8 and *j* > 8. The *P*-values obtained for miRNA and genes were 1.3 x 10^−4^ and 1.2 x 10^−11^, respectively. To detect a correlation between 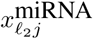 and 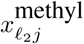 for *l*_2_ = 2, we constructed a scatter plot from these data (Fig. 1). It is obvious that they strongly correlate. *P*_*i*_ and *P*_*k*_ were computed by means of *x*_*l*_1_*i*_ and *x*_*l*_3_*k*_ of *l*_1_ = *l*_3_ = 2. Finally, we found seven miRNAs and 241 genes that showed differences in expression and promoter methylation, respectively, between control and treated samples. To confirm that the expression levels of seven miRNAs and the degrees of promoter methylation of 241 genes are correlated, we computed pairwise coefficients of correlation between them and computed adjusted *P*-values for each of these 7 × 241 = 1687 pairs. As presented in Table I, most of the pairs are associated with adjusted *P*-values less than 0.01, i.e., indicating statistical significance. Thus, TD-based unsupervised FE successfully identified pairs of miRNAs and genes having significant pairwise correlations.

**Figure 1.**
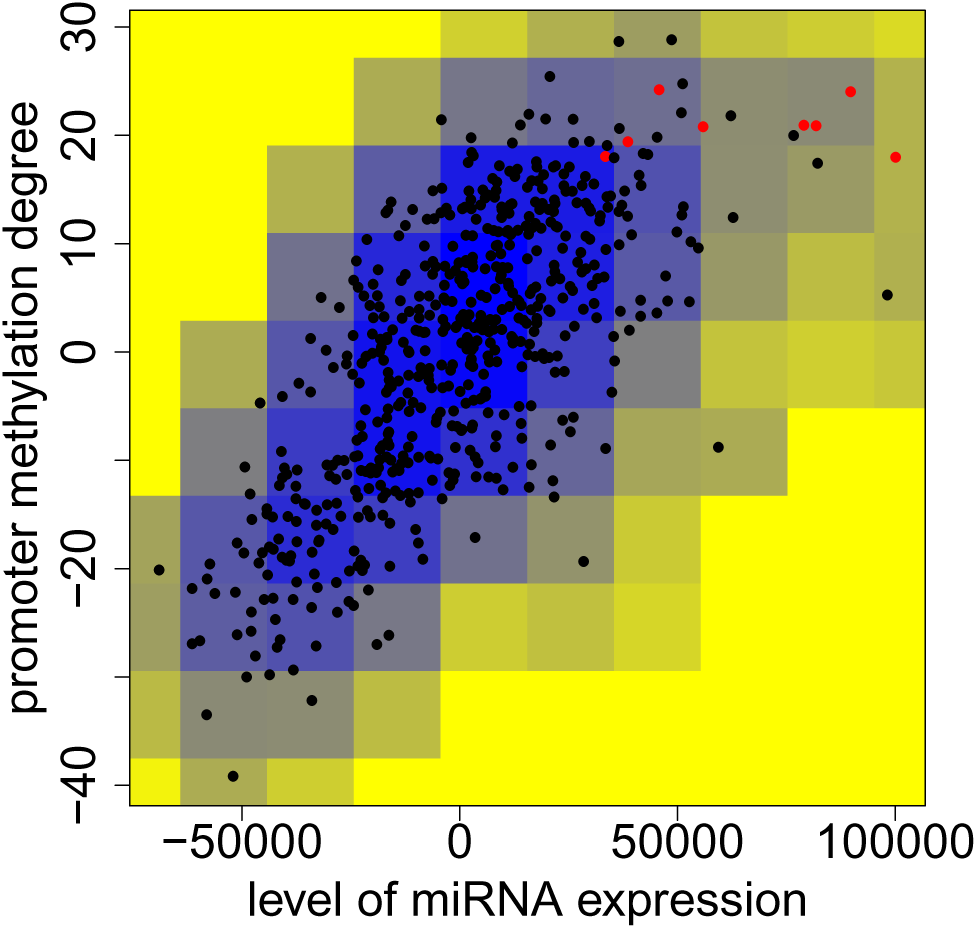
Scatter plot of 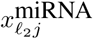 and 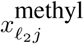. The coecient of correlation between them is 0.72 (*P* = 2.0 x 10^−92^). Red points and black points correspond to normal ovarian tissues and tumors, respectively. Color indicates local density of points (blue to yellow denote denser to sparser). All units are arbitrary.

**Table I.**
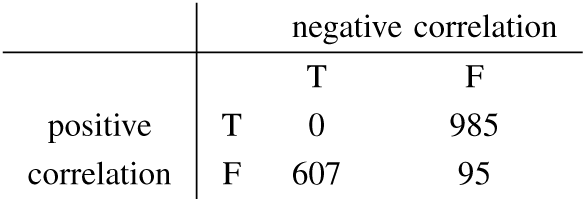
THE NUMBER OF MIRNA–GENE PAIRS SHOWING A SIGNIFICANT CORRELATION (ADJUSTED *P*-VALES LESS THAN 0.01) WHEN THEY ARE IDENTIFIED WITH TD-BASED UNSUPERVISED FE.

In order to see if other conventional methods can compete with TD-based unsupervised FE, we performed simple Student’s *t* test on miRNA expression data and degrees of promoter methylation of the genes. This analysis resulted in 214 miRNAs and 19395 genes associated with adjusted *P*-values less than 0.01 (by the BH criterion). This finding suggests that simple Student’s *t* test cannot select a reasonable number of miRNAs or genes associated with significant *P*-values. Next, to confirm the superiority of TD-based unsupervised FE toward simple Student’s *t* test, we computed pairwise coefficients of correlation between these 214 miRNAs and 19395 genes (Table II). In contrast to Table I where the majority of pairs show significant intrapair correlations, only ~13% pairs show a significant correlation. Thus, Student’s *t* test cannot identify pairs of miRNAs and genes with significant intrapair correlation with a small number of false positives. One may still wonder if Student’s *t* test can compete with TD-based unsupervised FE when correlation analysis is restricted to the miRNAs and genes with large differences between controls and treated samples. For this purpose, we repeated correlation analysis with top-ranked seven miRNAs and 241 genes according to *P*-values computed by Student’s *t* test (Table III). It is obvious that restricting the analysis to only top-ranked miRNAs and genes does not improve the identification of pairs with a significant correlation at all if Table III is compared with Table I.

**Table II.**
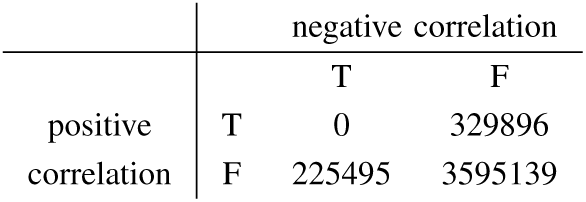
THE NUMBER OF MIRNA–GENE PAIRS SHOWING A SIGNIFICANT CORRELATION (ADJUSTED *P*-VALES LESS THAN 0.01) WHEN THESE ARE IDENTIFIED BY STUDENT–S *t* TEST.

**Table III.** THE NUMBER OF MIRNA–GENE PAIRS SHOWING A SIGNIFICANT INTRAPAIR CORRELATION (ADJUSTED *P*-VALES LESS THAN 0.01) WHEN ONLY SEVEN TOP-RANKED MIRNAS AND 241 TOP-RANKED GENES SELECTED BY STUDENT’S *t* TEST ARE CONSIDERED.

|  | negative correlation |  |  |
| --- | --- | --- | --- |
|  |  | T | F |
| positive correlation | T | 0 | 13 |
|  | F | 28 | 1646 |

Readers may still wonder whether selecting pairs with significant correlations prior to the identification of those showing a difference between controls and treated samples can select miRNAs and genes effectively. To test this idea, we computed *P*-values for all pairs of miRNA expression levels and the degrees of promoter methylation of genes and identified pairs with significant intrapair correlations (Table IV). Apparently, this approach is successful because only a limited number of pairs (less than 10%) were identified. Nevertheless, it cannot be used for identification of the miRNAs and genes with desired properties because it turned out that all the miRNAs and genes had at least one significant correlation.

**Table IV.** THE NUMBER OF MIRNA–GENE PAIRS (AN EXPRESSION LEVEL AND DEGREE OF PROMOTER METHYLATION, RESPECTIVELY) SHOWING A SIGNIFICANT INTRAPAIR CORRELATION (ADJUSTED *P*-VALES LESS THAN 0.01).

|  | negative correlation |  |  |
| --- | --- | --- | --- |
|  |  | T | F |
| positive correlation | T | 0 | 608989 |
|  | F | 588783 | 16809266 |

The above result suggests that TD-based unsupervised FE can outperform the conventional methods when selecting miRNAs and genes satisfying the three conditions presented in the Introduction section.

Next, we wanted to evaluate biological significance of the selected miRNAs and genes. First, seven miRNAs (Table V) were uploaded to DIANA-miRPath [16] with TarBase specified as a target identification database. Although a total of 66 KEGG pathways are enriched among these miRNAs (Table VI), there are at least 15 pathways related to cancer directly (bold). Next, gene symbols (Table V) were uploaded to MSigDB (http://software.broadinstitute.org/gsea/msigdb/annotate.jsp). C6: oncogenic signatures were tested and as many as 33 oncogenic expressed gene sets were found to significantly overlap (Only top twenty sets are included in Table VII because of lack of space).

**Table V.**
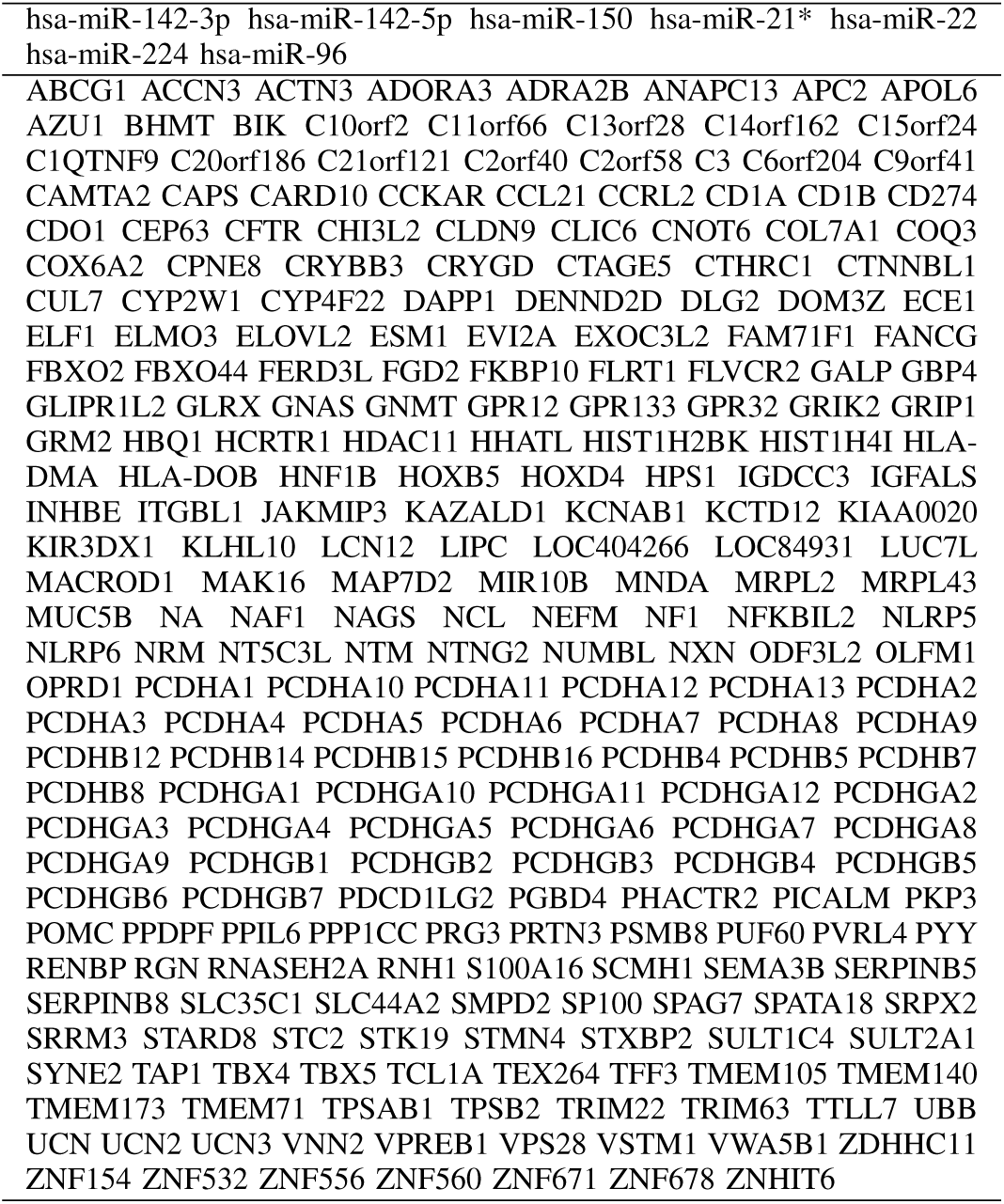
DNA-METHYLATION-REGULATED MIRNAS (7) AND GENES (241).

**Table VI.**
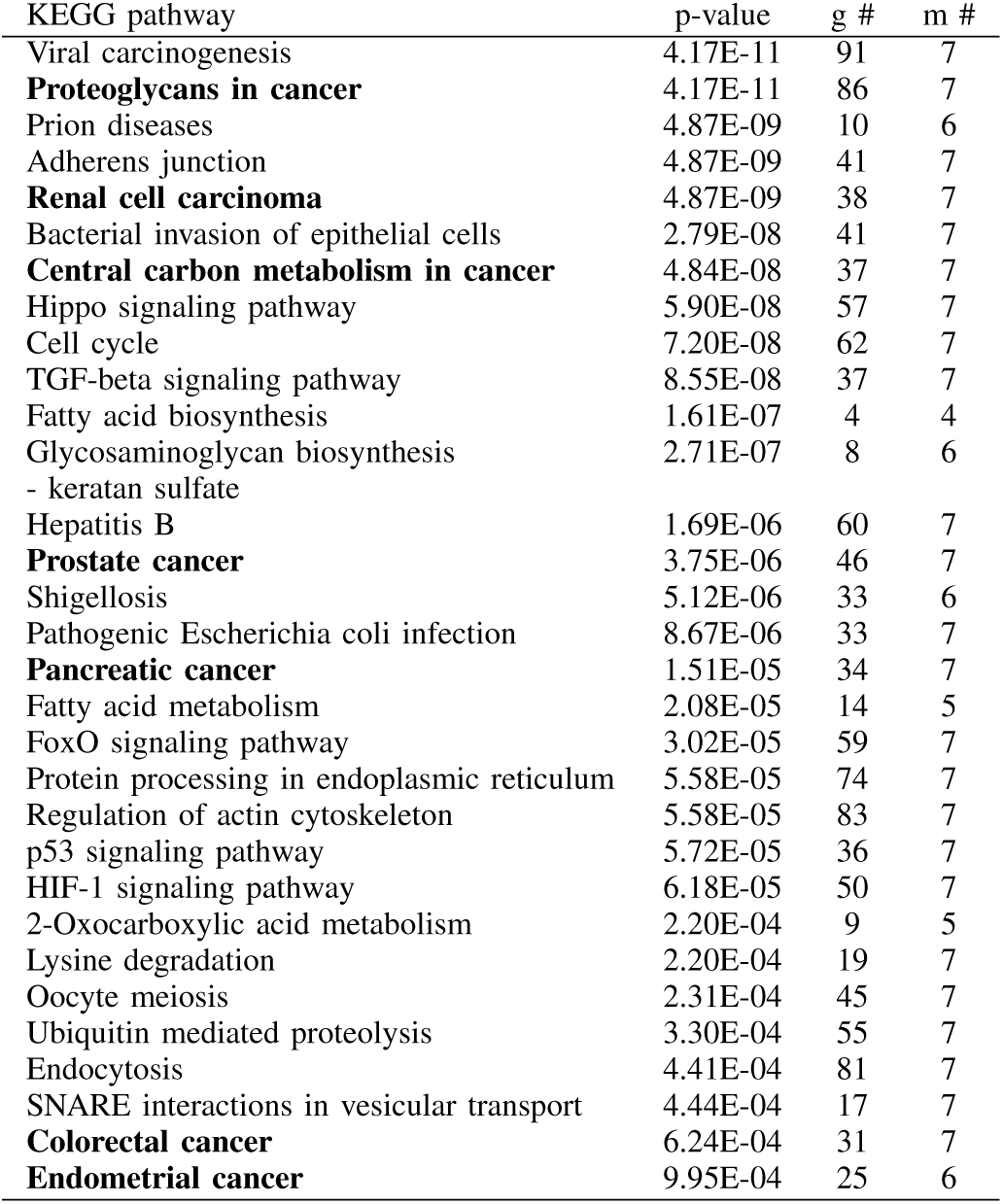
ENRICHED KEGG PATHWAYS DETECTED BY DIANA-MIRPATH AMONG SEVEN MIRNAS (TABLE V). BOLD ONES ARE CANCER RELATED. G #: THE NUMBER OF GENES, M #: THE NUMBER OF RELATED MIRNAS; *p*-VALUES ARE ADJUSTED.

**Table VII.** OVERLAPS BETWEEN C6: ONCOGENIC SIGNATURES IN MSIGDB AND GENE SYMBOLS (TABLE V) ASSOCIATED WITH GENES IDENTIFIED BY TD-BASED UNSUPERVISED FE AS SHOWING DIFFERENTIAL PROMOTER METHYLATION BETWEEN NORMAL OVARIAN TISSUES AND TUMOR TISSUES. #G1 (K): THE NUMBER OF GENES IN EACH OVEREXPRESSED GENES SET. #G2 (K): OVERLAPS WITH GENES SELECTED BY TD-BASED UNSUPERVISED FE.

| Gene Set Name | #G1 (K) | Description | # G2 (k) | k/K | p-value | FDR q-value |
| --- | --- | --- | --- | --- | --- | --- |
| WNT_UP.V1_DN | 170 | Genes downregulated in C57MG cells (mammary epithelium) by overexpression of WNT1 [Gene ID=7471] gene. | 9 | 0.0529 | 3.63E-07 | 4.58E-05 |
| KRAS.600.LUNG.BREAST_UP.V1_UP | 288 | Genes upregulated in epithelial lung and breast cancer cell lines overexpressing an oncogenic form of KRAS [Gene ID=3845] gene. | 11 | 0.0382 | 4.85E-07 | 4.58E-05 |
| RPS14_DN.V1_UP | 192 | Genes upregulated in CD34+ hematopoietic progenitor cells after a knockdown of RPS14 [Gene ID=6208] by RNA interference (RNAi). | 9 | 0.0469 | 1.01E-06 | 5.64E-05 |
| VEGF_A_UP.V1_UP | 196 | Genes upregulated in HUVEC cells (endothelium) by treatment with VEGFA [Gene ID=7422]. | 9 | 0.0459 | 1.19E-06 | 5.64E-05 |
| MEL18_DN.V1_UP | 141 | Genes upregulated in DAOY cells (medulloblastoma) upon a knockdown of PCGF2 [Gene ID=7703] by RNAi. | 7 | 0.0496 | 1.13E-05 | 4.26E-04 |
| KRAS.LUNG.BREAST_UP.V1_UP | 145 | Genes upregulated in epithelial lung and breast cancer cell lines overexpressing an oncogenic form of KRAS [Gene ID=3845] gene. | 6 | 0.0414 | 1.32E-04 | 3.37E-03 |
| KRAS.300_UP.V1_UP | 146 | Genes upregulated in four lineages of epithelial cell lines overexpressing an oncogenic form of KRAS [Gene ID=3845] gene. | 6 | 0.0411 | 1.37E-04 | 3.37E-03 |
| BMI1_DN.V1_UP | 147 | Genes upregulated in DAOY cells (medulloblastoma) upon a knockdown of BMI1 [Gene ID=648] by RNAi. | 6 | 0.0408 | 1.43E-04 | 3.37E-03 |
| KRAS.600_UP.V1_UP | 287 | Genes upregulated in four lineages of epithelial cell lines overexpressing an oncogenic form of KRAS [Gene ID=3845] gene. | 8 | 0.0279 | 1.64E-04 | 3.44E-03 |
| CAHOY_ASTROGLIAL | 100 | Genes upregulated in astroglia cells. | 5 | 0.05 | 2.02E-04 | 3.82E-03 |
| ATF2_UP.V1_DN | 187 | Genes downregulated in myometrial cells overexpressing ATF2 [Gene ID=1386] gene. | 6 | 0.0321 | 5.19E-04 | 7.71E-03 |
| CYCLIN_D1_UP.V1_UP | 188 | Genes upregulated in MCF-7 cells (breast cancer) overexpressing CCND1 [Gene ID=595] gene. | 6 | 0.0319 | 5.33E-04 | 7.71E-03 |
| SRC_UP.V1_UP | 188 | Genes upregulated in primary epithelial breast cancer cell culture overexpressing SRC [Gene ID=6714] gene. | 6 | 0.0319 | 5.33E-04 | 7.71E-03 |
| PTEN_DN.V1_UP | 191 | Genes upregulated upon a knockdown of PTEN [Gene ID=5728] by RNAi. | 6 | 0.0314 | 5.80E-04 | 7.71E-03 |
| MTOR_UP.N4.V1_DN | 193 | Genes downregulated in CEM-C1 cells (T-CLL) by rapamycin (sirolimus) [PubChem = 6610346], an mTOR pathway inhibitor. | 6 | 0.0311 | 6.12E-04 | 7.71E-03 |
| KRAS.LUNG_UP.V1_UP | 141 | Genes upregulated in epithelial lung cancer cell lines overexpressing an oncogenic form of KRAS [Gene ID=3845] gene. | 5 | 0.0355 | 9.74E-04 | 1.15E-02 |
| ALK_DN.V1_UP | 145 | Genes upregulated in DAOY cells (medulloblastoma) after a knockdown of ALK [Gene ID=238] by RNAi. | 5 | 0.0345 | 1.10E-03 | 1.16E-02 |
| BMI1_DN_MEL18_DN.V1_UP | 145 | Genes upregulated in DAOY cells (medulloblastoma) upon a knockdown of BMI1 and PCGF2 [Gene ID=648, 7703] by RNAi. | 5 | 0.0345 | 1.10E-03 | 1.16E-02 |
| P53_DN.V2_UP | 148 | Genes upregulated in HEK293 cells (kidney fibroblasts) upon a knockdown of TP53 [Gene ID=7157] by RNAi. | 5 | 0.0338 | 1.21E-03 | 1.20E-02 |
| KRAS.50_UP.V1_UP | 48 | Genes upregulated in four lineages of epithelial cell lines overexpressing an oncogenic form of KRAS [Gene ID=3845] gene. | 3 | 0.0625 | 2.14E-03 | 2.03E-02 |

It is known that the majority of ovarian cancers is derived from the ovarian surface epithelium [17]. It is evident from Table VII, among the first ten MSigDB records, five have “epithelium cell” descriptions, which accounted for 50% of the records. For instance, some of the oncogenic signatures were found in epithelial cells, the gene set names are WNT_UP.V1_DN, KRAS.600.LUNG.BREAST_UP.V1 UP, KRAS.LUNG.BREAST_UP.V1 UP, KRAS.300 UP.V1 UP, and KRAS.600 UP.V1 UP. Furthermore, clinical studies suggest that two hormones, estrogen and progesterone, are involved in ovarian cancer formation [18]. Table VIII lists the top six Gene Ontology (GO) molecular functional annotations of genes returned by MSigDB. Two of the molecular-function records are hormone related: “hormone activity” and “peptide hormone receptor binding”; the results were what we expected. In summary, the gene sets we identified were in line with the cell type and hormone records in the enrichment analysis.

**Table VIII.**
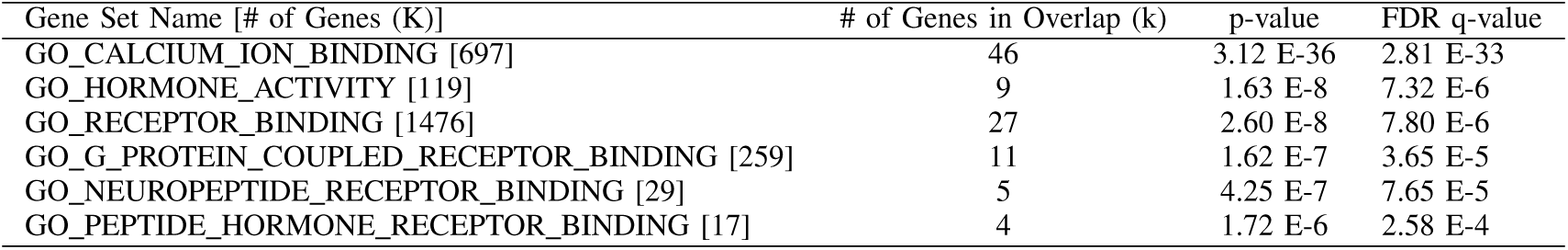
THE TOP SIX GO MOLECULAR FUNCTIONAL ANNOTATIONS OF THE GENES.

Tables VI, VII, and VIII suggest that TD-based unsupervised FE successfully identified cancer-related miRNAs and genes as expected.

## IV. CONCLUSION

In this paper, we applied TD-based unsupervised FE to miRNA expression and gene promoter methylation data (on ovarian tumors) retrieved from TCGA. TD-based unsupervised FE successfully identified genes with differential promoter methylation and differentially expressed miRNAs between normal ovarian tissues and tumors as well as significant correlations between the expression levels and methylation data. Student’s *t* test failed to identify the sets of miRNAs and genes satisfying these criteria. Biological evaluation of the identified miRNAs by DIANA-miRPath and of genes by MSigDB suggests that TD-based unsuper-vised FE identified genes and miRNAs related to cancers as expected.

ENRICHED KEGG PATHWAYS DETECTED BY DIANA-MIRPATH AMONG SEVEN MIRNAS (TABLE V). BOLD ONES ARE CANCER RELATED. G #: THE NUMBER OF GENES, M #: THE NUMBER OF RELATED MIRNAS; *p*-VALUES ARE ADJUSTED.

## ACKNOWLEDGMENT

This study was supported by KAKENHI 17K00417. Dr. Ka-Lok Ngs work is supported by the Ministry of Science and Technology (MOST) under the grants of MOST 106-2221-E-468-017, MOST 106-2632-E-468-002, and also supported by the grant from Asia University, 106-asia-06.

